# Environmental Stability of Enveloped Viruses is Impacted by the Initial Volume and Evaporation Kinetics of Droplets

**DOI:** 10.1101/2022.07.26.501658

**Authors:** Andrea J. French, Alexandra K. Longest, Jin Pan, Peter J. Vikesland, Nisha K. Duggal, Seema S. Lakdawala, Linsey C. Marr

## Abstract

Efficient spread of respiratory viruses requires the virus to maintain infectivity in the environment. Environmental stability of viruses can be influenced by many factors, including temperature and humidity. Our study measured the impact of initial droplet volume (50, 5, and 1 µL) and relative humidity (RH: 40%, 65%, and 85%) on the stability of influenza A virus, bacteriophage, Phi6, a common surrogate for enveloped viruses, and SARS-CoV-2 under a limited set of conditions. Our data suggest that the drying time required for the droplets to reach quasi-equilibrium (i.e. a plateau in mass) varied with RH and initial droplet volume. The macroscale physical characteristics of the droplets at quasi-equilibrium varied with RH but not with initial droplet volume. We observed more rapid virus decay when the droplets were still wet and undergoing evaporation, and slower decay after the droplets had dried. Initial droplet volume had a major effect on virus viability over the first few hours; whereby the decay rate of influenza virus was faster in smaller droplets. In general, influenza virus and SARS-CoV-2 decayed similarly. Overall, this study suggests that virus decay in media is closely correlated with the extent of droplet evaporation, which is controlled by RH. Taken together, these data suggest that decay of different viruses is more similar at higher RH and in smaller droplets and is distinct at lower RH and in larger droplets. Importantly, accurate assessment of transmission risk requires use of physiologically relevant droplet volumes and careful consideration of the use of surrogates.

**Funding:** National Institute of Allergy and Infectious Diseases, National Institute of Neurological Disorders and Stroke, National Institutes of Health; Department of Health and Human Services; Flu Lab.

**Importance:** During the COVID-19 pandemic, policy decisions were being driven by virus stability experiments involving SARS-CoV-2 applied to surfaces in large droplets at various humidity conditions. The results of our study indicate that determination of half-lives for emerging pathogens in large droplets likely over-estimates transmission risk for contaminated surfaces, as occurred during the COVID-19 pandemic. Our study implicates the need for the use of physiologically relevant droplet sizes with use of relevant surrogates in addition to what is already known about the importance of physiologically relevant media for risk assessment of future emerging pathogens.

## Introduction

Respiratory viruses, such as influenza A virus and SARS-CoV-2, contribute to high morbidity and mortality. These viruses must remain infectious in the environment for transmission to the next host to succeed. Understanding how environmental, host, and virus factors impact the stability of expelled virus will lead to a better assessment of virus transmission risk and ways to reduce it.

Many factors can impact virus stability in the environment, including virion structure, temperature, relative humidity (RH), droplet composition, solute concentration, and fomite surface material.^1–5^ However, the relationship between droplet size and virus stability is not well understood, even though droplet size plays an important role in virus transmission by governing the distance traveled by respiratory expulsions.^6^ Smaller droplets, or aerosols, can travel further from the infected host, while larger droplets settle to the ground more quickly due to their increased mass.^6^ Droplet or aerosol size can also be a determinant of host infection site, as those smaller than 10 μm in diameter are more likely to deposit deeper in the respiratory tract.^7^ Understanding how droplet volume affects virus stability is critical to mitigating transmission of respiratory viruses such as influenza virus and coronaviruses.

Studies measuring virus stability in the environment typically use one of two methods to produce the inoculum: nebulizers to produce aerosols, or pipettes to create droplets of volumes ranging from 5 to 50 μL. While large droplets are commonly used to assess environmental virus stability, they do not mimic a physiological volume of a droplet created by an expulsion. The vast majority of expelled droplets from the respiratory tract are less than 0·5 µL in volume (approximately 1 mm in diameter for a sphere). In contrast, a droplet of 50 μL (approximately 4·6 mm in diameter if spherical) is about 5 times larger and 100 times greater in volume.^8^ Studies measuring the stability of SARS-CoV-2 on surfaces have examined the virus in 5,^9^ 10,^10^ 20,^11^ or 50 µL droplets, ^2,3,12^ which are much larger than naturally expelled droplets. These initial studies into SARS-CoV-2 stability were used widely to assert the importance of fomite transmission and set policies. However, little work has been done to understand whether virus decay in large droplets is representative of decay in smaller, more physiologically relevant droplet volumes.

This study primarily used the 2009 pandemic influenza H1N1 virus (H1N1pdm09, A/CA/07/2009) and bacteriophage Phi6, a commonly used virus surrogate, to examine environmental stability of enveloped viruses in three different droplet volumes at three different RHs over time. Specifically, we measured the viability of each virus in 50, 5, and 1 µL droplets on surfaces over time at 40%, 65%, and 85% RH. We observed that virus within smaller droplets decays quickly regardless of RH, while virus decay occurs more slowly in larger droplets. We also explored droplet evaporation rates and found that virus decay is closely correlated with the extent of evaporation, which is likely a proxy for solute concentrations in the droplet. Additionally, limited experiments with SARS-CoV-2 showed that influenza virus decayed similarly to SARS-CoV-2 at an intermediate 55-60% RH in 50, 5, and 1 µL droplets. Overall, our results suggest that virus stability studies should use smaller, more physiologically relevant droplet volumes and should recognize the limitations of surrogate viruses.

## Results

### Relative humidity alters morphology of evaporating droplets and drying kinetics

We expect the physical and chemical characteristics of droplets to influence decay of viruses within each droplet. Some of these characteristics may be reflected in the morphology of droplets after they have dried.^5^ Furthermore, fluid dynamics within droplets could lead to increased aggregation of virus, which can enhance virus stability.^13,14^ We investigated whether droplet morphology and drying pattern at 24 hours differed between 1×50, 5×5, and 10×1 µL droplets (i.e., one droplet of volume 50 µL, five droplets of volume 5 µL, ten droplets of volume 1 µL). Droplets of Dulbecco’s Modified Eagle Medium (DMEM) were placed on polystyrene plastic and incubated at 40%, 65%, or 85% RH for 24 hours (Figure 1). These RH were selected to match values in other published work.^2^ At 40% RH, feather-like crystals grew throughout the dried droplet. At 65% RH, we observed fewer distinct crystals in the interior of the dried droplet. At 85% RH, the droplets retained a sheen, and crystallization did not occur. The effect of RH on dried droplet morphology was independent of initial droplet volume. The morphology of droplets containing virus was the same as that of droplets consisting of media alone (data not shown). Our results show that the droplet drying pattern at 24 hours depends on RH but not initial droplet volume, so any differences in viral decay by droplet size would not be due to final physico-chemical differences.

**Figure 1.**
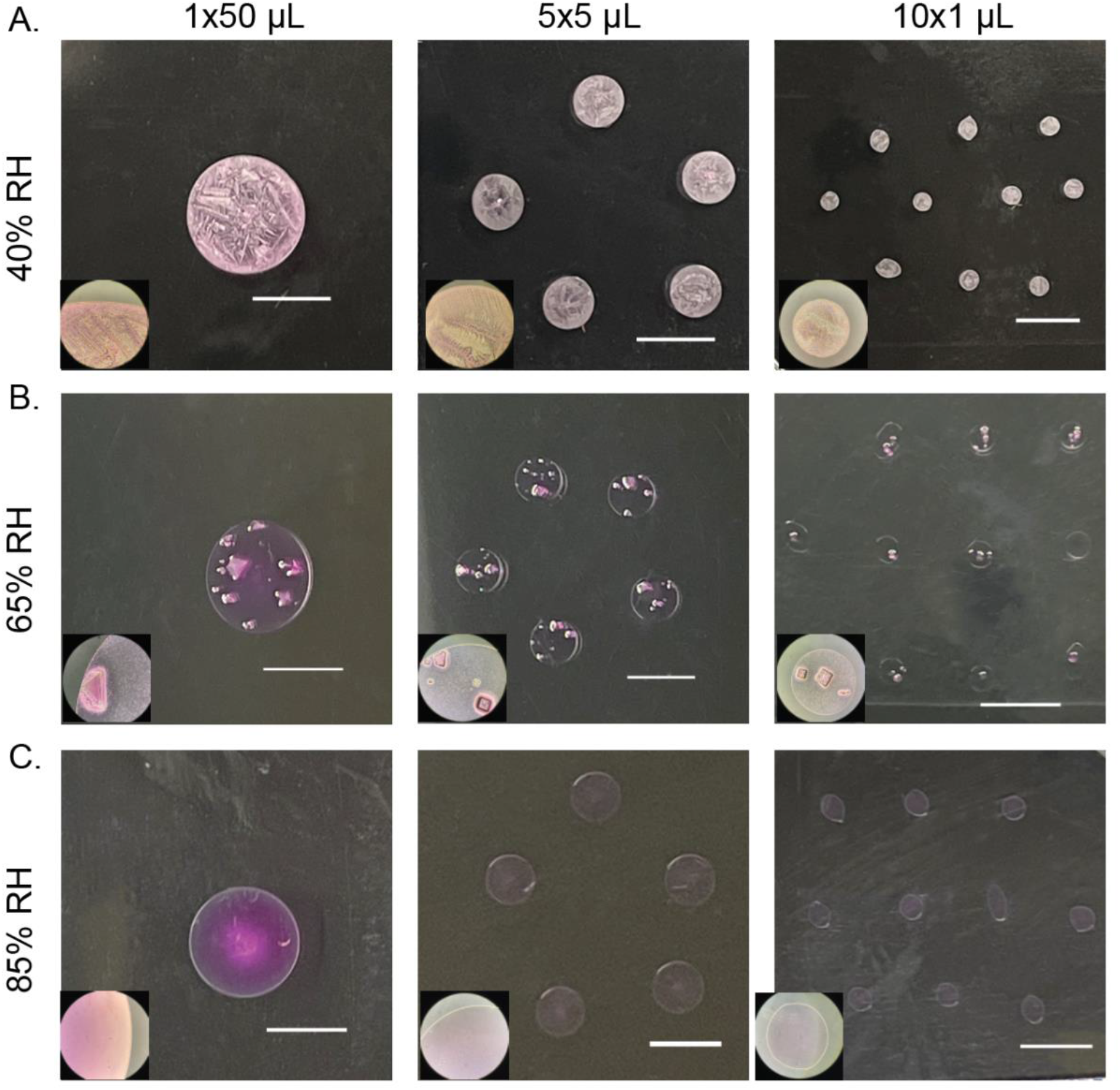
Macroscale physico-chemical characteristics of DMEM droplets vary with RH but not initial volume. Inset images taken with 10x objective. Scale bars indicate 5 mm. **A**. At 40% RH, droplets become concentrated at the border and develop interior feather-like crystals. **B**. At 65% RH, droplets develop distinct crystals within the interior. **C**. At 85% RH, droplets maintain moisture and do not crystallize.

Evaporation leads to the concentration of solutes, which can influence virus stability in droplets.^15^ To investigate the drying kinetics, we recorded the mass of droplets over time for 24 hours at ambient temperature and the same three RHs. At all RHs, the droplets lost mass linearly over time before reaching a plateau, referred to as a quasi-equilibrium (Supplemental Figure 1, Supplemental Table 1).^2^ We defined this state as quasi-equilibrium because it is likely that very slow evaporation continues over a much longer time scale until complete dryness occurs, or until a crust or shell forms that blocks further water loss. To simplify discussion, we refer to the time period before this as the “wet” phase and the period after this as the “dry” phase. Evaporation was faster for smaller droplets and lower RH (Supplemental Table 2). The time to reach quasi-equilibrium ranged from 0·5 hours for 1 µL droplets at 40% RH to 11 hours for 50 µL droplets at 85% RH. These data indicate that droplets of different volumes undergo different drying kinetics. If the kinetics of drying affect virus stability, then it could differ by initial droplet volume.

### Virus decay is more sensitive to relative humidity in large droplets

To directly examine how RH and droplet volume impact virus stability, we applied virus in droplets of different volumes to a polystyrene surface and quantified recovery of infectious virus over time.^16^ We compared decay of H1N1pdm09 and Phi6 at in 1×50 µL, 5×5 µL, and 10×1 µL droplets at 40%, 65%, and 85% RH (Figure 2). In 50 µL droplets, Phi6 decayed fastest at 40% RH and slowest at 85% RH (Figure 2A). The impact of RH on the decay of H1N1pdm09 in 50 µL droplets over the first 8 hours was similar as for Phi6 but less pronounced, with the fastest decay occurring at 40% RH (Figure 2B). Decay of H1N1pdm09 in the 50 µL droplet was first detected at 4 hours at 40% RH, 8 hours at 65% RH, and 24 hours at 85% RH, indicating that early virus decay was inversely related to RH (i.e., faster decay at lower RH) (Supplemental Table 4). Decay of Phi6 in 5 µL droplets differed by RH only at 1 hour, when decay at 40% was greater than at 85%. In 1 µL droplets, decay differed between 40 minutes and 4 hours by RH but was not significantly different at 8 hours and afterward (Figure 2A). H1N1pdm09 in 1 µL and 5 µL droplets decayed at a similar rate regardless of RH (Figure 2B). Phi6 was more unstable after drying at the intermediate RHs, whereas H1N1pdm09 tended to be more stable. This accounts for differences in decay at the smaller droplet sizes. These findings show that the impact of RH on virus decay in droplets depends on the virus and the initial volume of the droplets.

**Figure 2.**
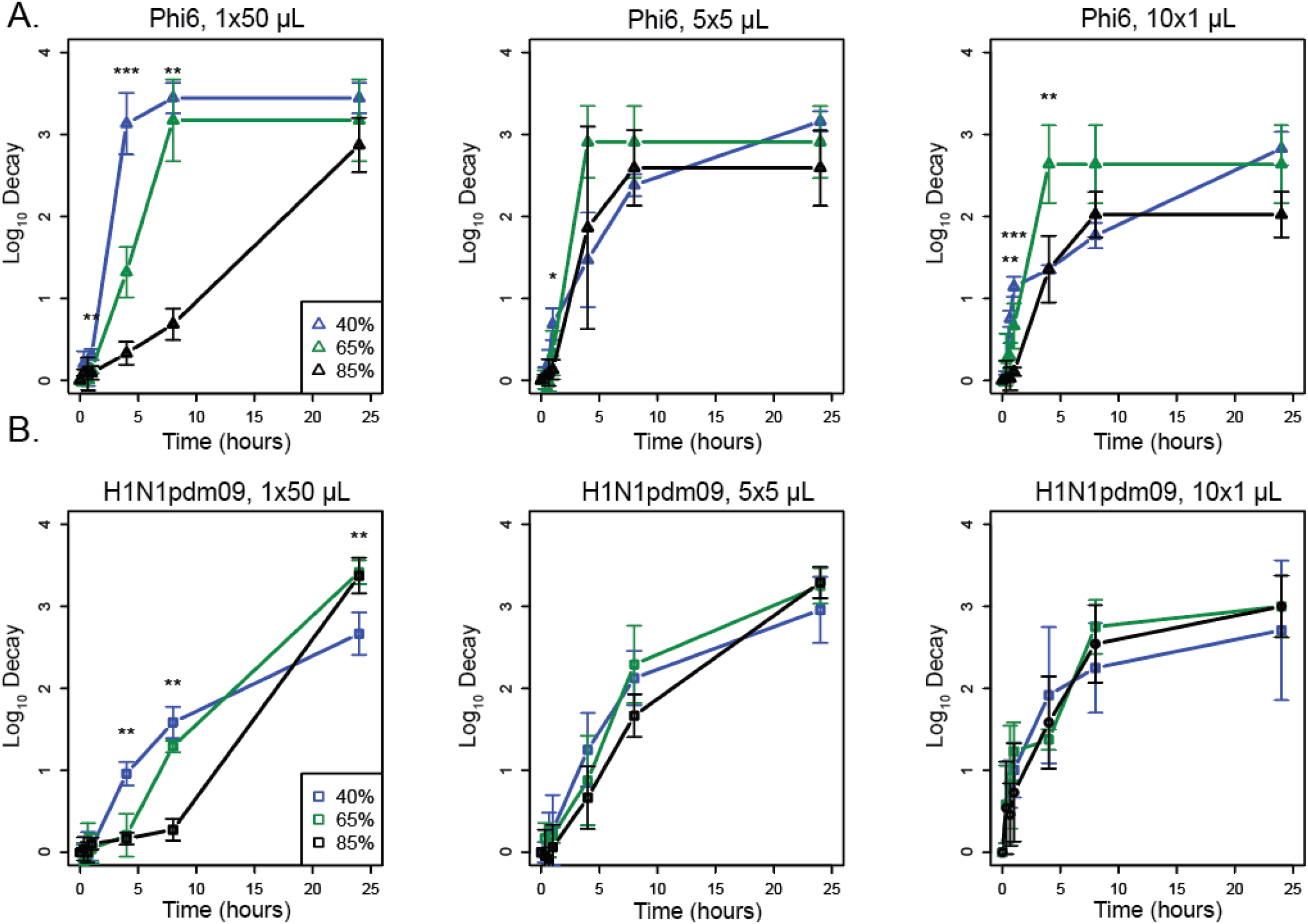
Virus decay varies more with relative humidity in large droplets than in small droplets. A. Titers of Phi6 in 1×50 µL, 5×5 µL, or 10×1 µL droplets compared at 40%, 65%, and 85% RH in terms of log_10_ decay. **B**. Titers of H1N1pdm09 in 1×50 µL, 5×5 µL, or 10×1 µL droplets compared at 40%, 65%, and 85% RH in terms of log_10_ decay. Error bars show standard deviation. Asterisks indicate significant differences between two or three RHs. For all graphs N=3 except at 1 hour where H1N1pdm09 N=6. One-way ANOVA tests were conducted between the RHs at each time point. A Tukey HSD test was conducted to determine between which RHs the significant differences (p <0·05) occurred. Statistical details can be found in Supplemental Table 3.

### Virus decay rates are faster during the wet phase and depend on droplet volume and virus

The pattern of decay for Phi6 and H1N1pdm09 appeared distinct for different droplet volumes. This led us to investigate whether evaporation rate impacts virus decay and whether different viruses behave similarly across different droplet volumes. Virus decay often follows first-order kinetics.^17^ Following a previously developed mechanistic model of virus inactivation in droplets, we fit an exponential decay model to virus titers in droplets during the wet phase (prior to quasi-equilibrium) and a separate curve during the dry phase to create a biphasic model (Figures 3 and 4, Table 1).^4^ The model accounts for changing solute concentrations in the droplets as they evaporate during the wet phase.^2^ In most cases, decay was faster in the wet phase than in the dry phase. Figure 3 shows viability as a function of time for two conditions: 5×5 µL droplets at 40% RH (Figure 3A-B) and 10×1 µL droplets at 65% RH (Figure 3C-D). The insets show the detail during the first 1·5 hours (Figure 3B, D), when the droplets transitioned from wet to dry. Similar trends are evident in most of the nine combinations of initial volume and RH for both viruses (Figure 4 and Table 1).

**Figure 3.**
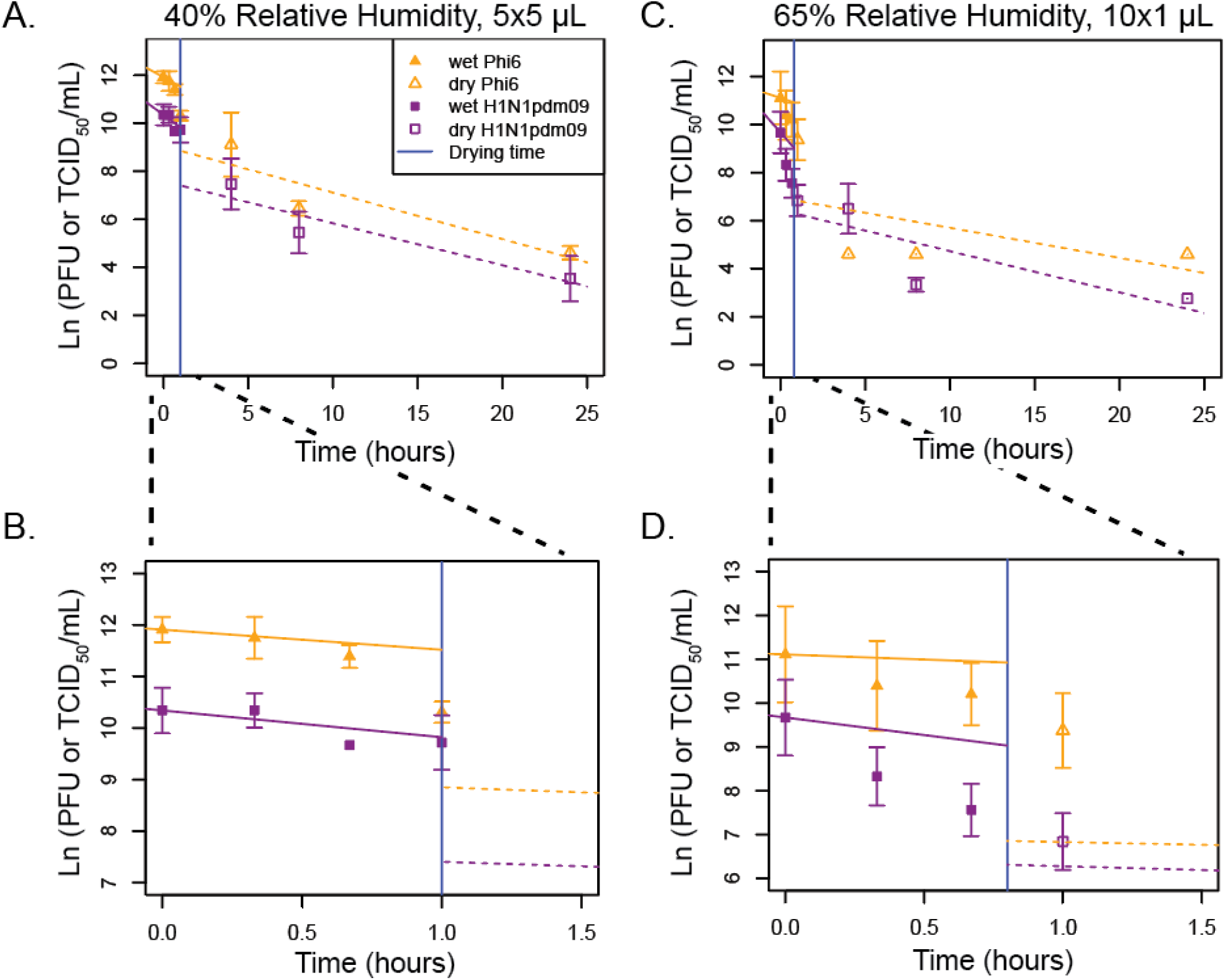
Mechanistic first-order decay modeling of viral decay in 5×5 µL droplets at 40% RH and 10×1 µL droplets at 65% RH shows that viral decay during the wet phase is greater than decay during the dry phase. **A-D**. First order exponential decay models, accounting for increasing solute concentrations over time during the wet phase, were fit to ln(PFU or TCID_50_/mL) over time for **(A-B)** 5×5 µL at 40% RH or **(C-D)** 10×1 µL droplets at 65% RH. **B**,**D** A magnification of A,C from 0 to 1·5 hours is shown. For all graphs N=3 except at 1 hour where H1N1pdm09 N=6. The vertical blue line indicates the time of transition from the wet phase to the dry phase. A t-test was used to compare the slopes between the evaporation and dry phases for each virus at each droplet volume and between the phases for each virus at each droplet volume (p <0·05). Statistical details can be found in Supplemental Table 5.

**Figure 4.**
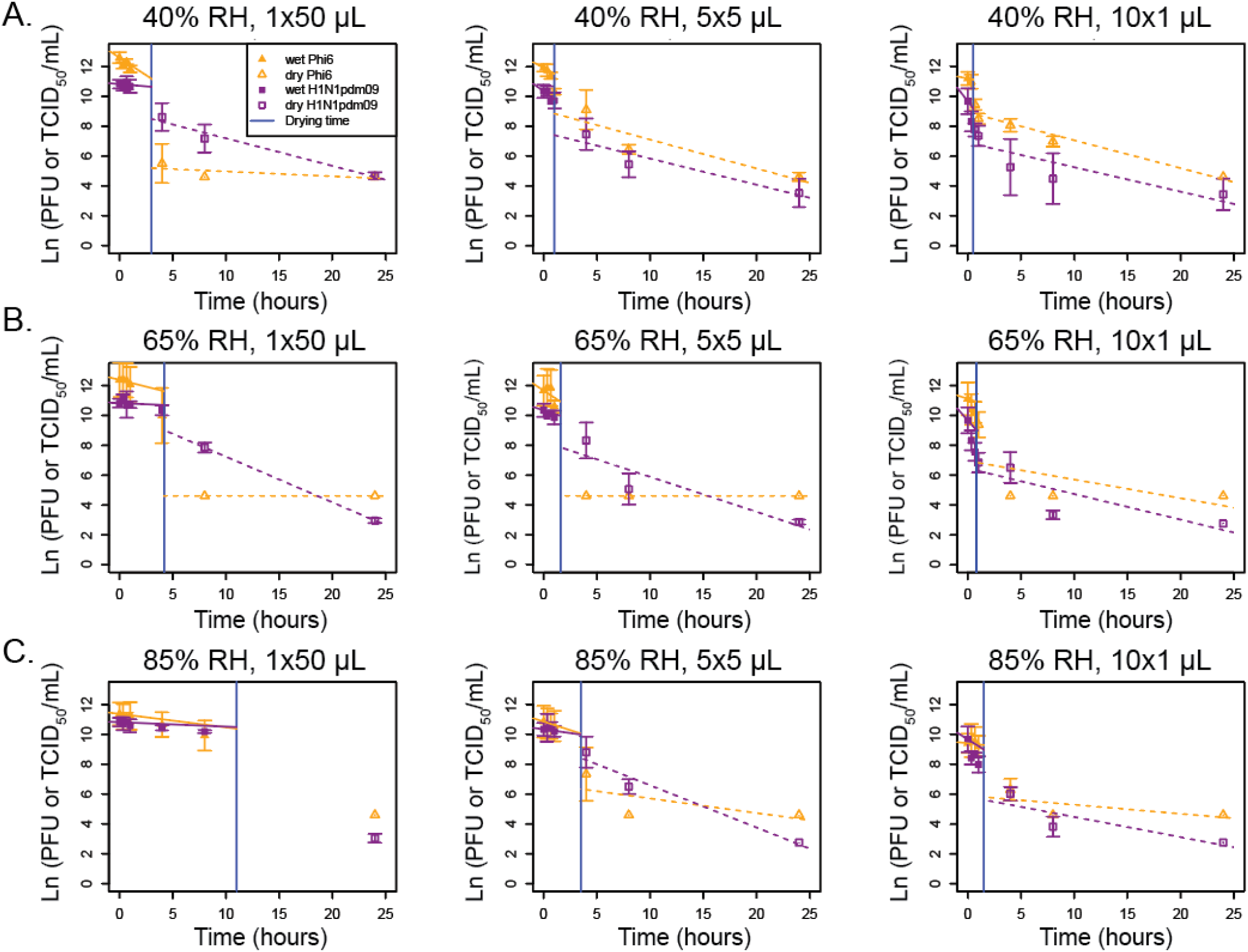
Mechanistic, first-order exponential decay modeling shows that the decay during the wet phase is greater or similar to the rate of decay the dry phase. **A-C**. First-order exponential decay models, accounting for increasing solute concentrations over time during the wet phase, were fit to ln(PFU or TCID_50_/mL) over time for 1×50 µL, 5×5 µL, or 10×1 µL droplets at **(A)** 40% RH, **(B)**, 65% RH, **(C)** or 85% RH. For the wet phase, the fitted initial decay rate is shown. For all graphs N=3 except at 1 hour where H1N1pdm09 N=6. The vertical blue line indicates the time of transition from the wet phase to the dry phase. A t-test was used to compare the slopes between the evaporation and dry phases for each virus at each droplet volume and between the phases for each virus at each droplet volume (p <0·05). Statistical details can be found in Supplemental Table 5.

**Table 1:**
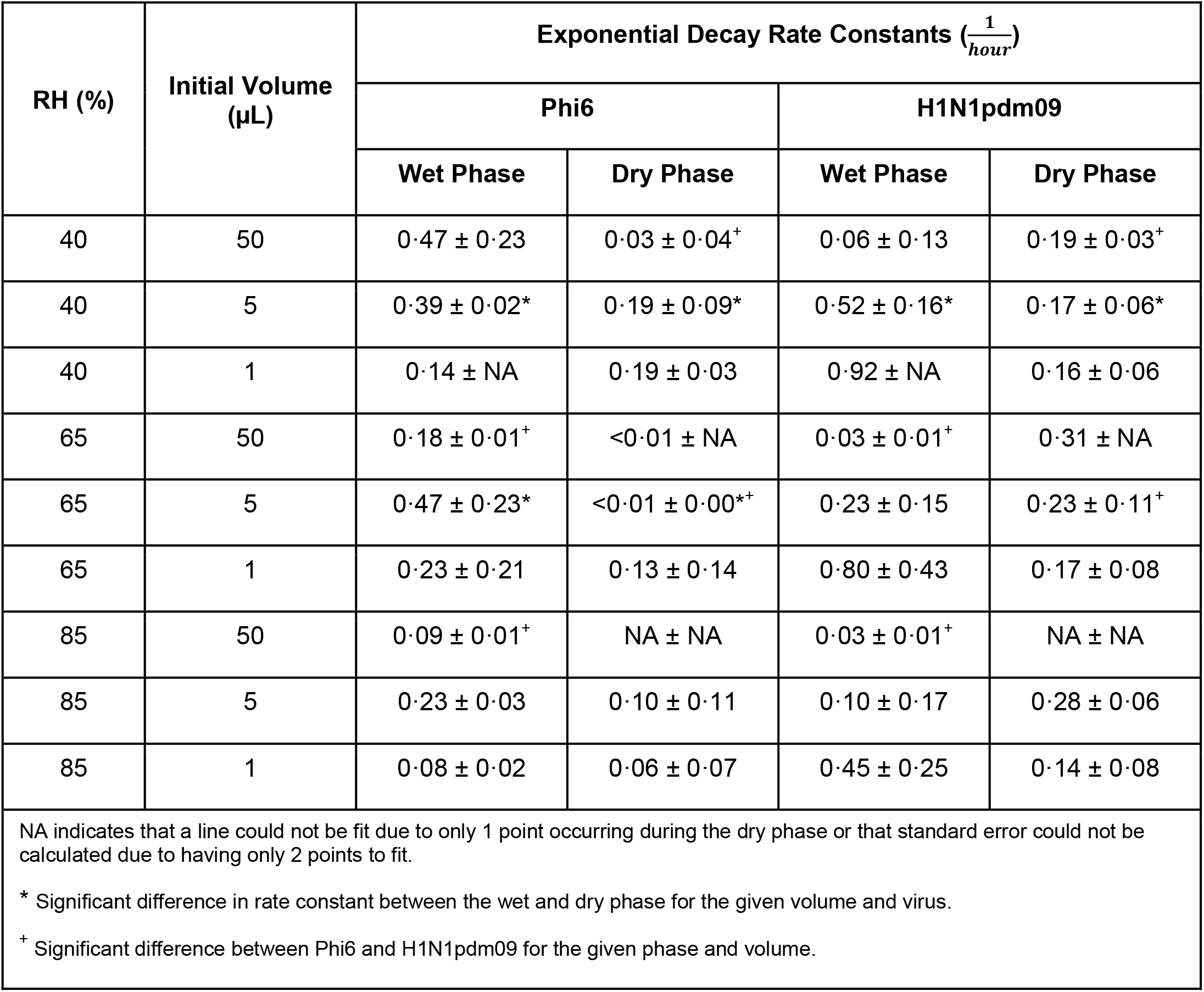
First-order exponential decay rate constants, adjusted for changing solute concentrations in the wet phase, for Phi6 and H1N1pdm09

Among the 12 combinations of RH and initial droplet volume for which the decay rate constant could be compared between the wet phase and dry phase, it was larger in magnitude during the wet phase than during the dry phase in most cases and significantly larger in 3 of 12 of these cases (Table 1, Supplemental Table 5): Phi6 in 5×5 µL droplets at 40% RH (Figure 3A), H1N1pdm09 in 5×5 µL droplets at 40% RH (Figure 3A), and Phi6 in 5×5 µL droplets at 65% RH (Figure 4B). Because there were only two time points during the wet phase for the 10×1 µL droplets at 40% RH and only one or two time points during the dry phase for 1×50 µL droplets at 65% and 85% RH, it was not possible to compare decay rates for these conditions (Table 1)

Comparing decay rates as a function of droplet size, we found that for H1N1pdm09 at a given RH and phase, the decay rate constant was often higher in smaller droplets compared to larger ones, mainly in the wet phase, although differences were only significant at 40% RH between 50 and 5 µL droplets in the wet phase. For Phi6, there were no apparent trends in the decay rate as a function of initial droplet volume.

Comparing decay rates by virus, we found that the decay rate constant was significantly higher for Phi6 than H1N1pdm09 in two cases during the wet phase and was significantly different in two cases—higher for H1N1pdm09 in both cases—during the dry phase (Figure 4, Table 1). Significant differences were not observed in the 1 µL droplets. These results indicate that different enveloped RNA viruses may decay differently.

### H1N1pdm09 decays similarly to SARS-CoV-2 at intermediate RH

Given the observed differences in the decay rate constants of H1N1pdm09 and Phi6 (Figure 4, Table 1), we further investigated how the stability of these two viruses compared to that of SARS-CoV-2 using both original and previously published data.^2^ To determine whether these viruses undergo similar patterns of decay at 40%, 65%, and 85% RH, we compared our results for H1N1pdm09 and Phi6 in 50 µL droplets to published results for SARS-CoV-2 (Figure 5, A-C, Supplemental Table 6).^2^ There were significant differences for each pairwise comparison of the decay of H1N1pdm09, Phi6, and SARS-CoV-2 at 40% RH at 4 and 8 hours. SARS-CoV-2 was most stable, followed by H1N1pdm09 and then Phi6 (Figure 5A). At 65% RH, there were fewer differences: only Phi6 was significantly different (less stable) from H1N1pdm09 and SARS-CoV- 2 again at 4 and 8 hours (Figure 5B). At 85% RH, there were no significant differences for the decay of any pairwise comparison (Figure 5C). Significance at 24 hours was not assessed due to virus decay reaching the limit of detection for at least one of the viruses tested. This suggests that in large 50 µL droplets, virus-specific differences are greater at lower RH.

**Figure 5.**
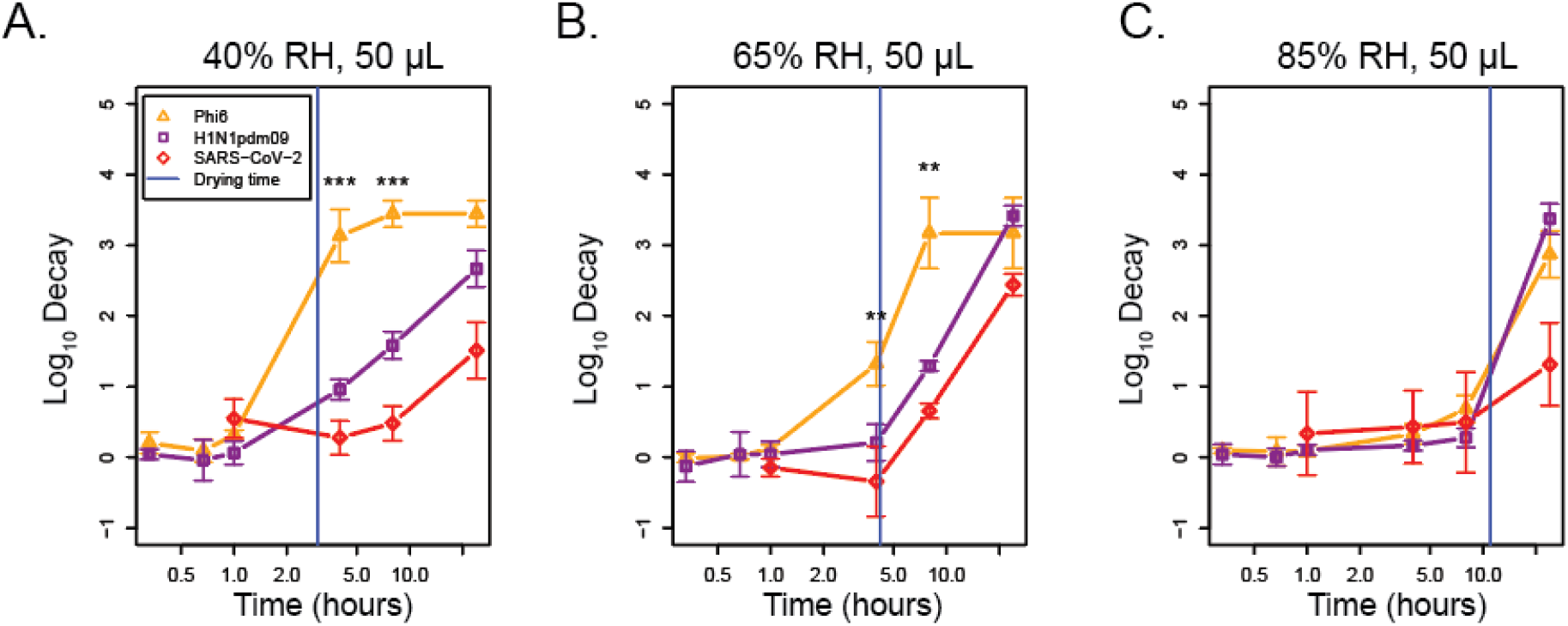
SARS-CoV-2 and H1N1pdm09 decay similarly across RH in 1×50 µL droplets. **A-C**. SARS-CoV- 2, H1N1pdm09, and Phi6 stability were measured at **(A)** 40%, **(B)** 65%, **(C)** and 85% RH in 50 µL droplets using SARS-CoV-2 data originally published in Morris et al.^2^ The vertical blue line indicates the time of transition from the wet phase to the dry phase. A one-way ANOVA and Tukey TSD test were used to determine statistical significance. Statistical details can be found in Supplemental Table 6.

To understand the role of droplet volume on decay of different enveloped respiratory viruses, we assessed titers of SARS-CoV-2 and H1N1pdm09 in 1×50 µL, 5×5 µL, and 10×1 µL droplets at 55% and 60% RH, respectively. Due to technical limitations, we were not able to test the exact same RH, but we consider these conditions to be similar. SARS-CoV-2 stability in 1×50 µL, 5×5 µL, and 10×1 µL droplets at 55% RH was similar to that of H1N1pdm09 at 60% RH, except for the 50 µL droplets at 4 hours (Figure 6, A-C, Supplemental Table 7). While decay of SARS-CoV-2 at 24 hours appeared to be greater, H1N1pdm09 had reached the maximum decay corresponding to the limit of detection. After 8 hours, decay was greatest in the 1 µL droplets and least in the 50 µL droplets. Taken together with Figure 5, these results show that SARS-CoV-2 and H1N1pdm09 decay similarly at intermediate RH and that differences in virus decay are evident in larger droplets.

**Figure 6.**
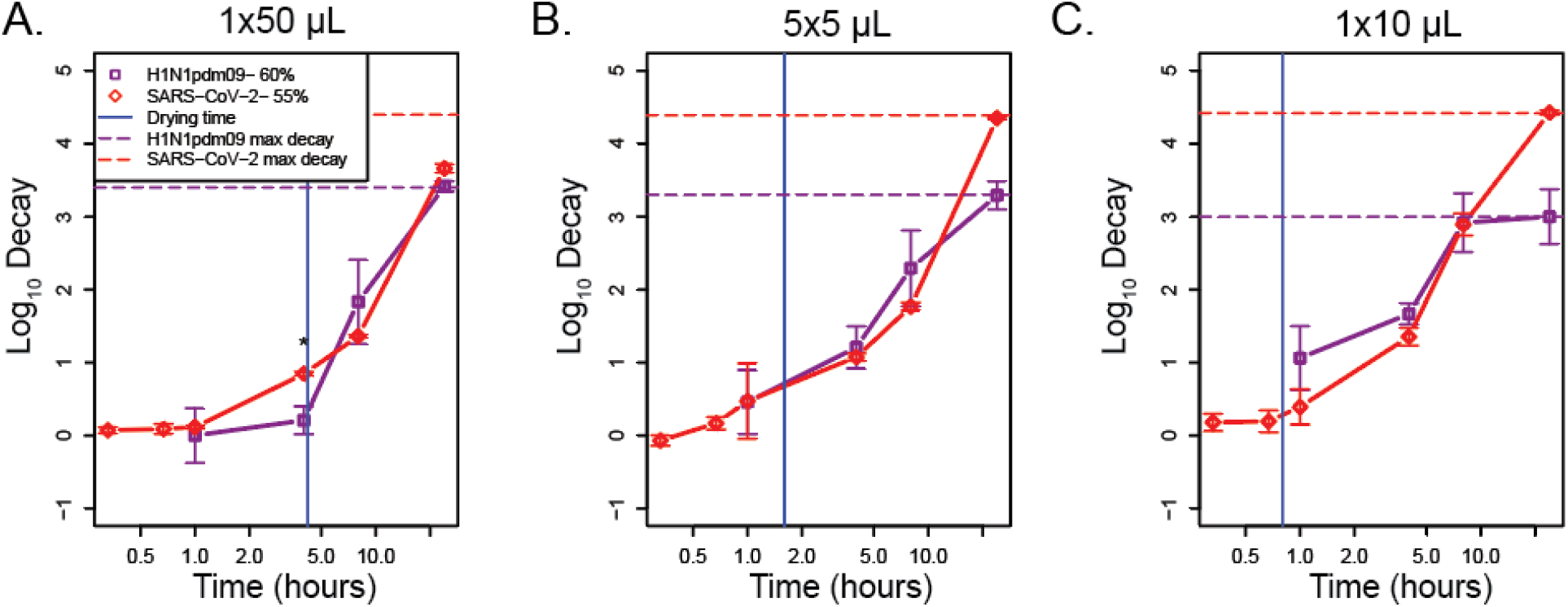
SARS-CoV-2 and H1N1pdm09 decay similarly across droplet volume at intermediate RH. **A-C**. SARS- CoV-2 stability was compared to H1N1pdm09 at 55-60% RH in 1×50 µL (**A**), 5×5 µL (**B**), and 10×1 µL (**C**) droplets over time. The vertical blue line indicates the time of transition from the wet phase to the dry phase. A one-way ANOVA was used to determine statistical significance. Statistical details can be found in Supplemental Table 7.

## Discussion

The studies detailed here characterize the interplay of droplet volume and RH on the stability of three enveloped RNA viruses: Phi6, H1N1pdm09, and SARS-CoV-2. Our results showed that RH has a greater impact on viral decay in large 50 µL droplets than in small 1 µL droplets, that decay rates during the wet phase are greater than or similar to decay rates during the dry phase regardless of droplet size and RH, and that differences in virus decay are more common in 50 µL droplets than in 1 µL droplets and at low RH.

Our results raise questions about the application of prior studies on stability of viruses that employ large droplet volumes to real-world transmission. For example, one study derived a half-life (∼0·3-log decay) of 6·8 hours for SARS-CoV-2 in 50 µL droplets on polypropylene plastic ^3^. Another used 5 µL droplets to evaluate the lifetime of SARS-CoV-2 on different materials, and reported 0·5-log decay in 3 hours and 1·1-log decay in 6 hours on plastic, similar to the results shown here (Figure 6B).^9^ The conclusions of studies might have differed had they used smaller, more physiologically relevant droplet volumes. In our study, after 4 hours, we observed no significant decay in 50 µL droplets, ∼1-log decay in 5 µL droplets, and ∼1.5-log decay in 1 µL droplets at 55-60% RH. Over longer time periods, results converged; we observed at least 3-log decay in all three droplet volumes after 24 hours. These differences are likely controlled by physical and chemical properties of the droplets as they undergo evaporation at different rates, depending on their initial volume and ambient humidity.

We have attempted to study viruses in a more realistic droplet volume compared to those used in past research, but even a 1 µL droplet is at the extremely large end of the range of droplet volumes observed in respiratory emissions. During talking, coughing, and sneezing, droplets of this size are emitted in much lower numbers, by many orders of magnitude, compared to those that behave as aerosols.^6^ While we observed differences between 50 µL and 1 µL droplets in this study, previous work has shown that virus in 1 µL droplets undergoes similar decay to that in aerosols at 23% to 98% RH and 22°C.^18^ New techniques are needed to study smaller droplet volumes on surfaces.

Our results also suggest caution in the use of surrogates to study the stability of pathogenic viruses and their potential for transmission.^19^ Surrogates can be useful for evaluating sampling and analysis methods, studying physico-chemical processes such as mechanisms of decay and transport in complex media, or eliciting trends in survival in complex media.^1,18,20,21^ However, we should be cautious about extrapolating survival times from surrogates to other viruses. In the present study, we found that Phi6 decayed more quickly than did H1N1pdm09 and SARS-Cov-2 under our experimental conditions. Relying on only Phi6 data could lead to incorrect, and potentially hazardous, conclusions about pathogenic viruses. Strain selection should also be considered when using influenza virus, as previous work has shown that avian influenza viruses undergo more rapid decay compared to human influenza viruses.^22^ Decay of enveloped viruses is likely dependent upon many complex interactions of media components with the viral glycoprotein and changes to it during and after drying. Thus variations in glycoprotein content and density per virus family or strain likely influence the stability within droplets. On the other hand, H1N1pdm09 decayed more similarly to SARS-CoV-2 and could be useful surrogate to extrapolate the latter’s persistence in more physiologically relevant conditions.

A major limitation of this study is that we used culture medium, DMEM, that may not be representative of real respiratory fluid. We selected this medium for the purpose of comparing results with prior studies of SARS-CoV-2 in DMEM.^2,3^ Prior studies have shown that virus survival in droplets, including suspended aerosols, is strongly dependent on the chemical composition of the suspending medium.^1,20,23^ In particular, we have previously shown that H1N1pdm09 in aerosols and 1 µL droplets survived better when the suspending medium was supplemented with extracellular material from human bronchial epithelial cells.^1^ Further studies will be required to characterize whether extracellular material from airway cells affects the decay patterns observed in this study.

Extrapolating our results to smaller droplet sizes and combining them with the findings of other studies may provide mechanistic insight into the dynamics of virus inactivation in droplets and aerosols. The biphasic virus decay that is readily observed in droplets likely occurs in aerosols, too, as shown in published work and a preprint.^2,9,24,25^ While a droplet/aerosol is wet and evaporation is still occurring, the virus is subject to a faster decay rate than after the droplet/aerosol reaches a solid or semi-solid state at quasi-equilibrium,^5^ as we also observed for all droplet sizes tested. At the point of efflorescence (the crystallization of salts as water evaporates), if it occurs, there appears to be a rapid loss in infectivity. With aerosols, the first phase occurs quickly, within seconds, and further observations of decay are dominated by the quasi-equilibrium phase. Thus, the first phase of decay is important for transmission at close range, when exposure occurs within seconds, whereas both phases are important for transmission at farther range.

Although virus stability in droplets and aerosols appears to be a complex function of droplet size, composition, humidity, and other variables, mechanistically their role is to modulate the microenvironment surrounding a virion, as suggested in published work and a preprint.^5,25,26^ Ultimately, molecular-scale interactions are what lead to virus inactivation. We combined results for all droplet sizes and all RHs and plotted virus decay as a function of extent of evaporation of a droplet, a proxy for its instantaneous physical and chemical characteristics, and found that the points appeared to converge more so than in Figures 2 and 4 (Supplemental Figure 2). The exact mechanisms of virus inactivation—the biochemical changes that occur—remain unknown and are ripe for further investigation.

Due to our findings on the sensitivity of virus persistence to both droplet volume and composition, we urge a shift toward the use of more realistic conditions in future studies. They should employ droplets as close in volume as possible to those released from the respiratory tract (sub-micron up to several hundred microns in diameter), and whose chemical composition closely mimics that of real respiratory fluid. These findings are critical for pandemic risk assessment of emerging pathogens and useful to improve public policy on optimal transmission mitigation strategies.

## Methods

### Evaporation experiment

Droplet mass was recorded every 10 minutes for up to 24 hours using a micro-balance. Experiments were performed in duplicate. Additional information can be found in the supplemental methods.

### Stability studies

We measured virus stability for Phi6 and H1N1pdm09 in a humidity-controlled chamber (Electro-Tech Systems) at room temperature and three (40%, 65%, and 85%) or four RHs, respectively (40%, 60%, 65%, and 85%). A logger (HOBO UX100-011) placed inside the chamber recorded relative humidity and temperature. Droplets were pipetted onto 6-well polystyrene tissue culture-coated plates (Thermo Scientific) in technical duplicates. Droplets were resuspended at seven different time points (0 minutes, 20 minutes, 40 minutes, 1 hour, 4 hours, 8 hours, and 24 hours), or four time points (1, 4, 8 and 24 hours) for the experiment at 60% RH, using 500 µL of DMEM containing 2% FBS, penicillin/streptomycin, and L-glutamine.

We measured the stability of SARS-CoV-2 in an airtight desiccator at room temperature and 55% RH as described previously.^18^ In short, we filled one polyethylene Petri dish with 10 to 20 mL of saturated magnesium nitrate solution and placed it and a fan at the bottom of the desiccator to control the humidity. After RH equilibrium was reached, usually within 5-10 minutes, we deposited droplets and later resuspended them as described previously. We measured virus titers by plaque assay on Vero cells. All collections were performed in technical duplicates and independent triplicates.

### Cells and viruses

Information regarding virus growth and quantification can be found in the supplement.

### Modeling

Virus decay calculations and modeling are described in the supplement.

## Contributors

SSL and LCM conceptualized and designed the study. NKD, SSL, and LCM supervised the study. AJF, AKL, and JP developed methodology, acquired data through plaque assays, and analyzed the data, and produced figures. PJV analyzed data on droplet characteristics. AJF and AKL wrote the first draft of the manuscript. All authors revised and edited the final version of the manuscript. All authors had access to all the data in the study and accept responsibility for the publication.

## Data sharing

The data collected in this study are publicly available in the Virginia Tech Data Repository at https://doi.org/10.7294/20134796.

## Acknowledgements

This work was supported by NIAID (CEIRS HHSN272201400007C, SSL and LCM), and in part with Federal funds from NIAID, NIH, and DHHS (CEIRR contract number 75N93021C00015, SSL). Additional funding was provided by Flu Lab (SSL and LCM), the Virginia Tech ICTAS Junior Faculty Award and NIH NINDS R01NS124204 (NKD). JF was supported by the University of Pittsburgh Training Program in Antimicrobial Resistance (T32AI138954). We thank Dr. Rachel Schwartz for critical review and feedback. We also thank Dr. Douglas Reed and his lab for allowing us access to the humidity cabinet used in these experiments.

